# Interactome Analysis Reveals Regulator of G Protein Signaling 14 (RGS14) is a Novel Calmodulin (CaM) Effector in Mouse Brain

**DOI:** 10.1101/247270

**Authors:** Paul R. Evans, Kyle J. Gerber, Eric B. Dammer, Duc M. Duong, Devrishi Goswami, Daniel J. Lustberg, Juan Zou, Jenny J. Yang, Serena M. Dudek, Patrick R. Griffin, Nicholas T. Seyfried, John R. Hepler

## Abstract

Regulator of G Protein Signaling 14 (RGS14) is a complex scaffolding protein with an unusual domain structure that allows it to integrate G protein and mitogen-activated protein kinase (MAPK) signaling pathways. RGS14 mRNA and protein are enriched in brain tissue of rodents and primates. In the adult mouse brain, RGS14 is predominantly expressed in postsynaptic dendrites and spines of hippocampal CA2 pyramidal neurons where it naturally inhibits synaptic plasticity and hippocampus-dependent learning and memory. However, the signaling proteins that RGS14 natively interacts with in neurons to regulate plasticity are unknown. Here, we show that RGS14 exists as a component of a high molecular weight protein complex in brain. To identify RGS14 neuronal interacting partners, endogenous RGS14 immunoprecipitated from mouse brain was subjected to mass spectrometry and proteomic analysis. We find that RGS14 interacts with key postsynaptic proteins that regulate neuronal plasticity. Gene ontology analysis reveals that the most enriched RGS14 interacting proteins have functional roles in actin-binding, calmodulin(CaM)-binding, and CaM-dependent protein kinase (CaMK) activity. We validate these proteomics findings using biochemical assays that identify interactions between RGS14 and two previously unknown binding partners: CaM and CaMKII. We report that RGS14 directly interacts with CaM in a calcium-dependent manner and is phosphorylated by CaMKII *in vitro*. Lastly, we detect that RGS14 associates with CaMKII and with CaM in hippocampal CA2 neurons by proximity ligation assays in mouse brain sections. Taken together, these findings demonstrate that RGS14 is a novel CaM effector and CaMKII phosphorylation substrate thereby providing new insight into cellular mechanisms by which RGS14 controls plasticity in CA2 neurons.

## Introduction

Synaptic plasticity is a vital process through which neurons modulate the strength of specific synaptic connections and is thought to encode key aspects of memory. Simultaneous neuronal activity and glutamate release at synapses activate intracellular and extracellular signaling cascades that induce plasticity. Regulator of G Protein Signaling 14 (RGS14) is a complex RGS protein that naturally suppresses synaptic plasticity in hippocampal CA2 neurons and hippocampus-dependent learning and memory ^1^. RGS14 mRNA and protein are highly expressed in the brains of adult rodents and primates ^2,3^. In the adult mouse brain, RGS14 predominantly localizes to CA2 pyramidal neurons in the hippocampus ^2^ where it is enriched postsynaptically in dendrites and spines ^1,3^. Unlike pyramidal neurons in the neighboring CA1 subregion, CA2 pyramidal neurons resist long-term potentiation (LTP) of synaptic transmission ^4^, a form of plasticity that is widely believed to serve as the cellular correlate of memory formation. Mice lacking RGS14 (RGS14 KO) display a robust and nascent capacity for LTP induction in CA2 neurons that is absent in wild-type (WT) littermates ^1^, and RGS14 KO mice also exhibit markedly enhanced spatial learning compared to WT littermates ^1^. Thus, RGS14 plays a critical role in naturally restricting CA2 synaptic plasticity and hippocampus-based learning.

RGS14 possesses multiple protein binding domains through which it engages various binding partners to integrate signaling pathways related to plasticity ^5,6^. Similar to other RGS proteins ^7^, RGS14 contains an RGS domain through which it directly binds activated Gαi/o-GTP subunits and stimulates their intrinsic GTPase activity to limit heterotrimeric G protein signaling ^8–10^. Unlike most other proteins, RGS14 contains a G protein regulatory (GPR) motif, an additional G protein interacting domain ^8,9,11^, through which it selectively binds inactive Gαi_1/3_-GDP to localize RGS14 to cellular membranes ^10,12–14^. RGS14 also contains two tandem Ras/Rap-binding domains (RBDs, R1 and R2) through which it preferentially interacts with activated H-Ras-GTP and Rap2-GTP as well as Raf kinases ^8,15–17^. RGS14 can functionally integrate G protein and MAPK signaling by inhibiting growth factor driven ERK1/2 activity through cooperative interactions between the GPR and first RBD ^16^. Consistent with this idea, the plasticity unmasked in CA2 neurons of RGS14 KO mice due to loss of RGS14 requires intact ERK activity ^1^. However, additional cellular mechanisms by which RGS14 governs plasticity in its host CA2 hippocampal neurons remain elusive.

While some studies have investigated RGS14 binding partners and signaling functions as recombinant proteins in ectopic systems^5,6^, none so far have examined these interactions in brain where RGS14 is enriched. Thus, to gain a better understanding of the signaling network RGS14 engages in CA2 neurons to limit synaptic plasticity therein, we initiated studies to identify endogenous binding partners of RGS14 in neurons. To accomplish this, we performed mass spectrometry analysis on native RGS14 protein complexes immunoprecipitated from mouse brain. We also validated two novel RGS14 interacting proteins identified in our proteomics screen using complementary biochemical and cellular approaches. Here we present the first characterization of the signaling interactome of native RGS14 in the mouse brain. We report that RGS14 exists in a large, multiprotein complex which includes numerous synaptic proteins that have been shown to influence plasticity induction. We next validate these protein interactions for two previously unknown RGS14 binding partners – calmodulin (CaM) and Ca^2+^/CaM-dependent protein kinase II (CaMKII) – using purified and recombinant protein assays. Lastly, we demonstrate that RGS14 interactions with CaM and CaMKII are detectable in mouse CA2 hippocampal neurons. These findings suggest new roles for RGS14 in regulating neuronal plasticity signaling and physiology in hippocampal area CA2.

## Experimental Procedures

### Plasmids and proteins

RGS14 and CaM were expressed and proteins purified as previously described ^10,18^. The rat RGS14 cDNA used in this study (GenBank accession number U92279) was acquired as described ^9^. FLAG-RGS14 truncation mutants containing residues 1-202, 205-490, 371-544, and 444-544 were created as described ^14^. The RGS14-Luciferase construct used in these studies was created as described ^19^. The FLAG-CaMKIIa plasmid was a generous gift from Chris Yun (Emory University School of Medicine).

### Animals

Adult male and female wild-type C57BL/6J mice were used in this study. Animals in all experiments were housed under a 12h:12h light/dark cycle with access to food and water *ad libitum*. All experimental procedures conform to US NIH guidelines and were approved by the animal care and use committees of Emory University and NIEHS/NIH.

### Immunoperoxidase staining and light microscopy

For immunohistochemical studies, adult wild-type C57BL/6J mice were deeply anesthetized with isoflurane and transcardially perfused with cold 0.9% saline solution, followed by 4% paraformaldehyde (PFA, w/v) in PBS, pH 7.4. After perfusion, the brains were removed from the skull and immersion fixed in 4% PFA in PBS, pH 7.4, for 24 hours at 4°C. Brains were embedded in paraffin and coronally sliced in 10 μm sections. Immunoperoxidase staining was performed as previously described ^2^ by incubating sections with an anti-RGS14 monoclonal antibody (NeuroMab, 1:500). Mouse brain sections were mounted and coverslipped on glass slides and analyzed using an Olympus B×51 light microscope.

### Synaptosome isolation

Synaptosomes were isolated from adult WT mouse brain as previously described ^20^. Briefly, brains from three adult WT mice were rapidly removed and homogenized, and cellular debris was pelleted by centrifugation at 1,000 × g for 10 mins at 4°C. Cleared supernatant was directly applied to a discontinuous Percoll gradient, and synaptosomes were recovered from the interface of 15 and 23% (vol/vol) Percoll layers and resuspended in isotonic buffer as described ^20^. Following SDS-PAGE, immunoblotting was performed with antibodies against RGS14 (NeuroMab, 1:300), PSD95 (Millipore, 1:5,000), and synaptophysin (Millipore, 1:25,000).

### Mouse brain co-immunoprecipitation and proteolytic digestion

Adult wild-type C57BL/6J mice were deeply anesthetized by isoflurane inhalation and euthanized by decapitation. Brains were rapidly removed from the skull and homogenized on ice using a glass dounce homogenizer with 10 strokes in an ice-cold buffer containing 50 mM Tris, 150 mM NaCl, 5 mM EDTA, 5 mM MgCl_2_, 2 mM DTT, Halt phosphatase inhibitor cocktail (1:100, Thermo Fisher), and one mini protease inhibitor cocktail tablet (Roche Applied Science), pH 7.4. Membranes were solubilized by the addition of 1% NP-40 for 1h at 4°C and subsequently centrifuged to pellet debris. Cleared brain homogenates were incubated with an anti-RGS14 mouse monoclonal antibody (5 μg, NeuroMab) for 2 h at 4°C. Next, 75 μl of Protein G Dynabeads (Thermo Fisher) were added to homogenates for 2 h to precipitate antibody-bound protein complexes. Protein G Dynabeads were washed thoroughly with ice-cold TBS and immediately digested for MS. IPs were simultaneously performed with generic mouse IgG (Millipore) as a control for comparative MS analysis. Three independent biological replicates were performed for each condition. Following four washes with ice cold TBS, the control beads or RGS14 immunoprecipitated samples were resuspended in 50 mM NH_4_HCO_3_ buffer and protein reduced with 1 mM dithiothreitol (DTT) at 25°C for 30 minutes, followed by 5 mM iodoacetimide (IAA) at 25°C for 30 minutes in the dark. Protein was then digested overnight with 12.5ng/μL trypsin (Promega) at 25°C. Resulting peptides were desalted with a stage tip and dried under vacuum.

### Mass spectrometry analysis of RGS14 interacting proteins

For LC-MS/MS analysis, peptides were resuspended in 10 μL of loading buffer (0.1% formic acid, 0.03% trifluoroacetic acid, 1% acetonitrile) essentially as previously described with slight modification ^21^. Peptide mixtures (2 μL) were separated on a self-packed C18 (1.9 μm Dr. Maisch, Germany) fused silica column (25 cm × 75 μM internal diameter (ID); New Objective, Woburn, MA) by a Dionex Ultimate 3000 RSLCNano and monitored on a Fusion mass spectrometer (ThermoFisher Scientific, San Jose, CA). Elution was performed over a 120 minute gradient at a rate of 250 nL/min with buffer B ranging from 3% to 80% (buffer A: 0.1% formic acid in water, buffer B: 0.1 % formic in acetonitrile). The mass spectrometer cycle was programmed to collect at the top speed for 3 second cycles. The MS scans (300-1500 m/z range, 200,000 AGC, 50 ms maximum ion time) were collected at a resolution of 120,000 at m/z 200 in profile mode and the HCD MS/MS spectra (1.5 m/z isolation width, 30% collision energy, 10,000 AGC target, 35 ms maximum ion time) were detected in the ion trap. Dynamic exclusion was set to exclude previous sequenced precursor ions for 20 seconds within a 10 ppm window. Precursor ions with +1, and +8 or higher charge states were excluded from sequencing.

### MaxQuant for label-free protein quantification

Raw data files were analyzed using MaxQuant v1.5.3.30 with Thermo Foundation 2.0 for RAW file reading capability. The search engine Andromeda was used to build and search a Uniprot mouse reference (downloaded on Aug 14, 2015). Protein Methionine oxidation (+15.9949 Da) and protein N-terminal acetylation (+42.0106 Da) were variable modifications (up to 5 allowed per peptide); cysteine was assigned a fixed carbamidomethyl modification (+57.0215 Da). Only fully tryptic peptides were considered with up to 2 miscleavages in the database search. A precursor mass tolerance of ±20 ppm was applied prior to mass accuracy calibration and ±4.5 ppm after internal MaxQuant calibration. Other search settings included a maximum peptide mass of 6,000 Da, a minimum peptide length of 7 residues, 0.5 Da Tolerance for ion trap HCD MS/MS scans. The false discovery rate (FDR) for peptide spectral matches and proteins were set to 1%. The label free quantitation (LFQ) algorithm in MaxQuant ^22,23^ was used for protein quantitation. LFQ intensity of each protein for each mouse was averaged from three bead control IP samples and three RGS14 IP samples. No more than two missing values were considered in the RGS14 IP samples, which were imputed as previously described ^24^. Differentially expressed proteins were found by calculating Student’s t-test p values and fold difference |log2(*RGS14*/*non-specific IgG*)| ≥ 0.58 (≥ ±1.50 fold change). Volcano plots were plotted with ggplot2 package in R. The mass spectrometry proteomics data have been deposited to the ProteomeXchange Consortium via the PRIDE ^25^ partner repository with the dataset identifier PXD008461.

### Gene Ontology enrichment analysis and visualization

Functional enrichment of the differentially aggregated proteins was determined using the GO-Elite (v1.2.5) package ^26^ as previously described ^27^. The set of total proteins identified and quantified (*n*=1,362) was used as the background. The input list included proteins significantly interacting (p<0.05) with RGS14 (*n*=233). Positive Z-score determines degree of over‐representation of ontologies and Fisher exact P-value was used to assess the significance of the Z-score. For both increased and decreased proteins in the insoluble proteome, a Z-score cut off of 1.96, (P-value cut off of 0.05) with a minimum of 3 proteins per category was employed. Horizontal bar graph was plotted in R. A visualization of the GO terms for the RGS14 interactors was performed using the EnrichmentMAP plugin for Cytoscape v.3.2.1, after converting GO‐Elite pruned output to DAVID format ^28,29^. A subset of proteins falling within six of those ontologies was further visualized as a circular chord plot using the R GOplot package.

### CaM-Agarose pull-down assays

For CaM-Agarose pull-down assays with purified RGS14 protein, 25 μl of CaM-Agarose beads (Sigma) were washed twice with a binding buffer composed of 20 mM HEPES, 150 mM NaCl, 0.1% Tween-20 (v/v), pH 7.5 supplemented with either 0.1 mM CaCl_2_ (“+ Ca^2+^”) or 5mM EDTA (“– Ca^2+^”). For immunoblotting, 0.25 μg of purified RGS14 protein was diluted in each binding buffer, and 10% of these samples were removed as input samples. The remaining RGS14 protein was incubated with pre-washed CaM-Agarose beads for 2 h at 4°C, and beads were then thoroughly washed in the appropriate binding buffer. Proteins were eluted with Laemmli buffer, heated at 95°C in a heating block, and subjected to SDS-PAGE and immunoblotting.

For CaM-Agarose pull-down assays with recombinant FLAG-RGS14 expressed in HeLa cells, cell lysates were prepared as described above. 50 μl of CaM-Agarose beads washed twice with a binding buffer composed of 20 mM HEPES, 100 mM NaCl, pH 7.5 supplemented with either 2 mM CaCl_2_ (“+ Ca^2+^”) or 5mM EDTA (“– Ca^2+^”). 10% of the initial cell lysates was removed as input samples for immunoblotting. The remaining RGS14 protein was incubated with pre-washed CaM-Agarose beads for 2 h at 4°C, and beads were then thoroughly washed in the appropriate binding buffer. Proteins were eluted with Laemmli buffer, heated at 95°C for 5 mins, and subjected to SDS-PAGE and immunoblotting.

### Dansyl-CaM fluorescence measurements

Steady-state fluorescence spectra were recorded using a QM1 fluorescence spectrophotometer (Photon Technology International) with a xenon short arc lamp at 25°C as previously described ^18^. For dansyl-CaM fluorescence measurement, 0.8 mL solution containing 0.5 μM dansyl-CaM in 10 mM HEPES, 100 mM KCl, pH 7.5 with 2 mM Ca^2+^ or 5 mM EGTA was titrated with 5–10 μL aliquots of the RGS14 protein in the same buffer. The fluorescence spectra were recorded between 400 and 600 nm with an excitation wavelength at 335 nm and the slit width set at 4–8 nm. Experiments were repeated three times to obtain the average value. The normalized fluorescence intensity changes were plotted as a function of RGS14 concentration.

### Cell culture and transfection

HeLa cells (ATCC) were maintained in Dulbecco’s modified Eagle’s medium (Mediatech) supplemented with 10% fetal bovine serum (Atlanta Biologicals, 5% after transfection), 2 mM L‐ glutamine (Invitrogen), 100 U/mL penicillin (Mediatech), and 100 mg/mL streptomycin (Mediatech) in a humidified environment at 37°C with 95%O_2_/5% CO_2_. Transfections were performed using previously described protocols with polyethyleneimine (PEI; Polysciences) ^30^. After 24 hours of expression, cells were washed with PBS and harvested in an ice-cold lysis buffer containing 50 mM Tris, 150 mM NaCl, 2 mM DTT, 5 mM MgCl_2_, 1% Triton X-100 (v/v), phosphatase inhibitors (1:1,000; Sigma Aldrich), and one mini protease inhibitor cocktail tablet (Roche Applied Science), pH 7.4. Lysis buffer was supplemented with 2 mM CaCl_2_ or 5 mM EDTA (“+ Ca^2+^” and “– Ca^2+^”, respectively). Cells were lysed for 1h at 4°C rotating end-over-end, and subsequently centrifuged to pellet cell debris. Cleared cell lysates were then subjected to CaM-Agarose pull-down assays or co-immunoprecipitation prior to immunoblotting.

### SDS-PAGE and immunoblotting

Samples were loaded onto 11% acrylamide gels and subjected to sodium dodecyl sulfate-polyacrylamide gel electrophoresis (SDS-PAGE) to separate proteins. Proteins were then transferred to nitrocellulose and subjected to immunoblotting to probe for proteins of interest. After blocking nitrocellulose membranes for 1 hour at room temperature in blocking buffer (5% nonfat milk (w/v), 0.1% Tween-20, and 0.02% sodium azide, diluted in 20 mM TBS, pH 7.6), membranes were incubated with primary antibodies diluted in the same buffer overnight at 4°C, except for anti-FLAG primary antibody. An anti-RGS14 mouse monoclonal antibody (NeuroMab) was used at a 1:1,000 or 1:5,000 dilution to detect recombinant or purified proteins, respectively. An anti-His mouse monoclonal antibody (Qiagen) was used at a dilution of 1:500 to detect purified H_6_-Gα_i1_. Membranes were washed in TBS containing 0.1% Tween-20 (TBST) and subsequently incubated with an anti-mouse (1:5,000) horseradish peroxidase (HRP)-conjugated secondary antibody diluted in TBST for 1 hour at room temperature. Following block, anti-FLAG-HRP (1:35,000, Sigma) primary antibody was diluted in TBST and incubated with membranes for 1 hour at room temperature with no secondary antibody. Protein bands were visualized using enhanced chemiluminescence and exposing membranes to films.

### Blue Native-PAGE

For blue native (BN)-PAGE, mouse brain lysates were separated on 4–16% Novex BisTris gels (Invitrogen) according to manufacturer’s instructions. After electrophoresis, gels were transferred to a PVDF membrane overnight, and the following day morning, membranes were washed in PBS containing 0.1% Tween 20 (PBST) for several hours to remove excess Coomassie blue. Immunoblotting was performed as described above using an anti-RGS14 mouse monoclonal antibody (NeuroMab, 1:300) to detect native protein complexes.

### Hydrogen/deuterium exchange (HDX) mass spectrometry

Solution phase amide HDX was carried out with a fully automated system as described previously ^31^. Briefly, 5 μl of 10 μM RGS14 was diluted to 25 μl with D_2_O-containing HDX buffer and incubated at 4°C for 10, 30, 60, 900, or 3,600 s. Following on exchange, back-exchange was minimized, and the protein was denatured by dilution to 50 μl in a low pH and low temperature buffer containing 0.1% (v/v) TFA in 3 M urea (held at 1°C). Samples were then passed across an immobilized pepsin column (prepared in house) at 50 μl min^−1^ (0.1% (v/v) TFA, 15°C); the resulting peptides were trapped on a C8 trap cartridge (Hypersil Gold, Thermo Fisher). Peptides were then gradient eluted from 4% (w/v) CH_3_CN to 40% (w/v) CH_3_CN, 0.3% (w/v) formic acid over 5 min at 2°C across a 1 × 50-mm C18 HPLC column (Hypersil Gold, Thermo Fisher) and electrosprayed directly into a Q Exactive hybrid quadrupole-Orbitrap mass spectrometer (Q Exactive, Thermo Fisher). The peptide set for the on-exchange experiments was generated from a separate tandem MS experiment where peptide ion signals were confirmed if they had a MASCOT score of 20 or greater and had no ambiguous hits using a decoy (reverse) sequence. The intensity-weighted average m/z value (centroid) of each peptide’s isotopic envelope was calculated with software developed in house ^32^ and corrected for back-exchange on an estimated 70% recovery and accounting for the known deuterium content of the on-exchange buffer. To measure the difference in exchange rates, we calculated the average percentage of deuterium uptake for RGS14 following 10, 30, 60, 900, and 3,600 s of on exchange. From this value, we subtracted the average percentage of deuterium uptake measured for the Ca^2+^/CaM:RGS14 complex (2:1 molar ratio).

### *In vitro* CaMKII phosphorylation assays

Purified CaMKII (NEB) was first pre-activated for 10 mins at 30°C in NEBuffer for protein kinases containing 50mM Tris-HCl, 10mM MgCl_2_, 0.1 mM EDTA, 2mM DTT, 0.01% Brij 35, pH 7.5, supplemented with 2 mM CaCl_2_, 1.2 μM CaM, and 200 μM ATP. 50 U of pre-activated CaMKIIα was then incubated for 20 mins at 30°C with 4×10^6^ cpm of [Υ −^32^P]-ATP (Perkin Elmer) and 2 μg of purified RGS14 or hexa-histidine-tagged Gαi1 (H_6_-Gα_i1_). A small amount of purified proteins (2.5%) were set aside as input samples for immunoblotting. Reactions were quenched by the addition of Laemmli sample buffer and heating at 95°C for 5 mins. Proteins were then separated by SDS-PAGE, and acrylamide gels were dried and exposed to film to detect phosphorylation by autoradiography.

### HeLa cell co-immunoprecipitation (co-IP)

RGS14-Luciferase and FLAG-CaMKIIα were co-transfected into HeLa cells and lysed in the presence/absence of Ca^2+^ as described above. 50 μl of anti-FLAG M2 agarose affinity gel (Sigma) or Protein G sepharose beads (GE, negative control) were pre-blocked with 3% BSA and then incubated with cell lysates for 3h at 4°C. Beads were washed thoroughly in ice cold tris-buffered saline (TBS), and proteins were eluted with an equal volume of Laemmli buffer, heated at 95°C for 5 mins, and subjected to SDS-PAGE and immunoblotting.

### Proximity Ligation Assays and Imaging

Adult wild-type C57BL/6J mice were deeply anesthetized with an overdose of sodium pentobarbital (250 mg/kg) and transcardially perfused with 4% paraformaldehyde in PBS. Brains were postfixed for 24 h, submerged in 30% sucrose in PBS, embedded in OCT (TissueTek), and sectioned coronally at 40 μm on a cryostat. RGS14 interactions with CaM and CaMKIα in hippocampus were detected using the Duolink In Situ Red Starter Kit for Mouse/Rabbit (Millipore Sigma) according to manufacturer instructions. Briefly, free floating brain sections containing hippocampus were subjected to antigen retrieval (3 min in 10 mM sodium citrate buffer at 100° C) followed by incubation in blocking solution for two hours at room temperatures. Sections were incubated overnight with primary antibodies against RGS14 (mouse, NeuroMab, 1:500) and either CaMKIIα (rabbit, Abcam, 1:250) or CaM (rabbit, SySy, 1:250). Proximity ligation assays were then carried out according to manufacturer instructions to produce red (624 nm) fluorescence when immunolabeled proteins are within 40 nm ^33^. Images were acquired on a Zeiss 710 meta confocal microscope using a 63X oil-immersion lens using consistent laser power and Z-stack parameters between samples. Images were processed using FIJI software (NIH v2.0.0). Images shown are representative results of PLAs performed in brain sections from three biological replicates. All images were only adjusted uniformly together as a group for brightness/contrast and then cropped for presentation.

## Results

Previous studies have reported that RGS14 protein is enriched in rodent ^1–3,9^ and primate ^3^ brain tissue. Immunostaining of adult mouse brain sections with a specific anti-RGS14 antibody shows RGS14 protein predominantly localizes to pyramidal neurons in hippocampal area CA2 (Fig 1A), which is consistent with our previous findings ^1–3^. Immunoblotting for endogenous RGS14 from WT mouse brain after SDS-PAGE detects a single band at the predicted molecular weight of ~61 kDa (Fig 1B, left). To investigate the protein complexes that RGS14 associates with in brain, mouse brain lysates were separated by BN-PAGE and immunoblotting was performed with the same anti-RGS14 antibody. Under non-denaturing conditions, RGS14 migrates as a series of protein bands ranging from ~150 – 450 kDa (Fig 1B, right), suggesting that native RGS14 exists as part of one or more large multi-protein complexes in mouse brain. Next, we isolated synaptic compartments from mouse brain by subcellular fractionation to determine if RGS14 is present at synapses as suggested by electron microscopy studies ^1,3^. We detect RGS14 protein is present in the Percoll gradient fraction containing synaptosomes and in recovered synaptosomes from mouse brain (Fig 1C).

**Figure 1.**
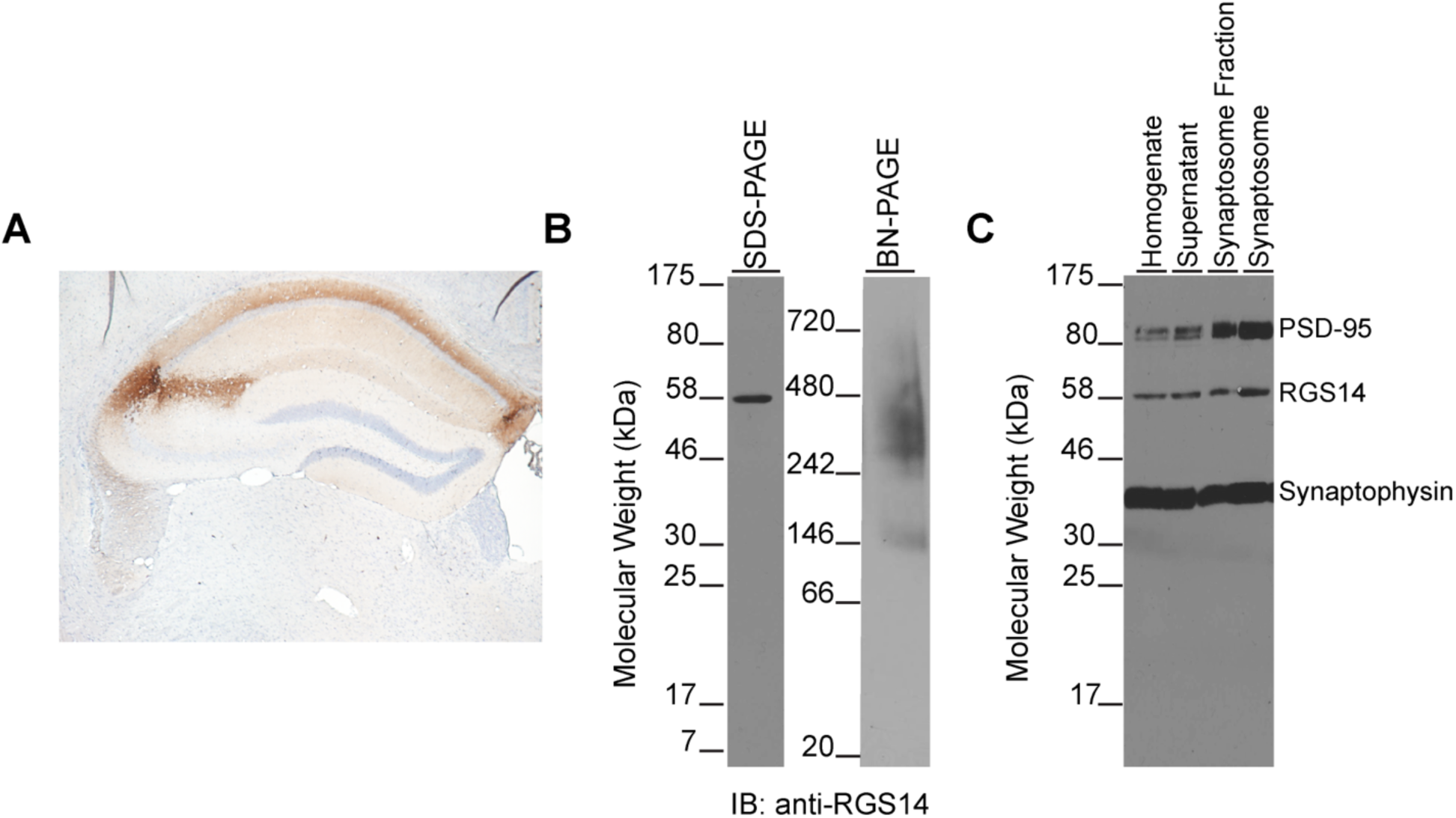
RGS14 exists as part of a high molecular weight protein complex in mouse brain. A) Immunoperoxidase labeling for RGS14 protein in adult mouse brain shows restricted expression to hippocampal CA2 neurons. Note: background deposition in CA1 neuropil reflects RGS14-positive CA2 axons that target area CA1. B) Immunoblots for endogenous RGS14 from mouse brain lysate separate by SDS-PAGE (left) and blue native gel electrophoresis (right). Under denaturing conditions, RGS14 runs near its predicted molecular weight (61 kDa), but blue native gel electrophoresis reveals native RGS14 from mouse brain migrates at much higher molecular weights. Results are representative of three replicates. C) Immunoblots of mouse brain synaptosome preparations reveal RGS14 is present in synaptic compartments. The molecular markers synaptophysin and PSD-95 indicate both pre‐ and post-synaptic elements, respectively, are present to compose a synaptosome. 100 μg of total protein was loaded in each lane; results are representative of four replicates.

To determine the proteins composing these large molecular weight assemblies in complex with RGS14, endogenous RGS14 was immunoprecipitated (IP) from mouse brain and subjected to mass spectrometry to identify interacting proteins as described in Experimental Procedures. The complete MS data are available via ProteomeXchange with the identifier PXD008461 (see also Tables S1 and S2). Of the 1,362 total proteins identified and quantified, 233 specific RGS14 interactors were significantly enriched over IgG IP controls. All 233 hits met the following explicit selection criteria: the IP and control IgG pulldown were performed in triplicate, LFQ intensity missing values for the RGS14 were controlled so that imputation could not drive the differences, and a T-test p value of <0.05 was enforced to arrive at the list of 233 candidate interactors. These data are shown in the half volcano plot (Fig 2A) with significantly enriched RGS14 interactors highlighted on the right boxed in yellow. A gene ontology (GO) analysis of RGS14 interacting proteins is shown in Figure 2B with hits grouped according to biological process, molecular function, and cellular component plotted as a function of Z-score. Of note, the most significant molecular function clusters represent actin binding, calmodulin(CaM)-dependent protein kinase (CaMK) activity, and CaM binding – all of which are critical for long-term potentiation (LTP) in the hippocampus. An enrichment map is provided for all GO hits (Fig 2C) that depicts the discrete GO terms and the number of genes that overlap between these groups. The top enriched RGS14 interacting proteins are displayed in the circle plot ordered by enrichment with corresponding GO terms on the right (Fig 3). Consistent with the postsynaptic localization of RGS14 in CA2 pyramidal neurons ^1^, functional annotation groups a substantial portion of the interacting proteins to dendritic spines and synapses. Of particular interest are the interactors corresponding to CaMK activity (red) as these signaling proteins have central roles in postsynaptic mechanisms of LTP ^34^ and all four isoforms of CaMKII were identified as RGS14 interactors. A critical step in LTP induction is postsynaptic Ca^2+^ influx through NMDA-type glutamate receptors, which then binds CaM (Ca^2+^/CaM) and drives the activity of numerous effectors, including CaMKII. The lack of plasticity in CA2 neurons has been attributed to more active Ca^2+^ buffering and extrusion in spines relative to CA3/CA1 ^35^, and these differences are due, at least in part, to the actions of the CaM-binding protein PCP4/Pep19. Given that RGS14 also suppresses LTP in CA2 neurons ^1^ and several proteins identified in our proteomics screen cluster into CaM binding (Fig 2B), we next investigated if RGS14 directly binds CaM.

**Figure 2.**
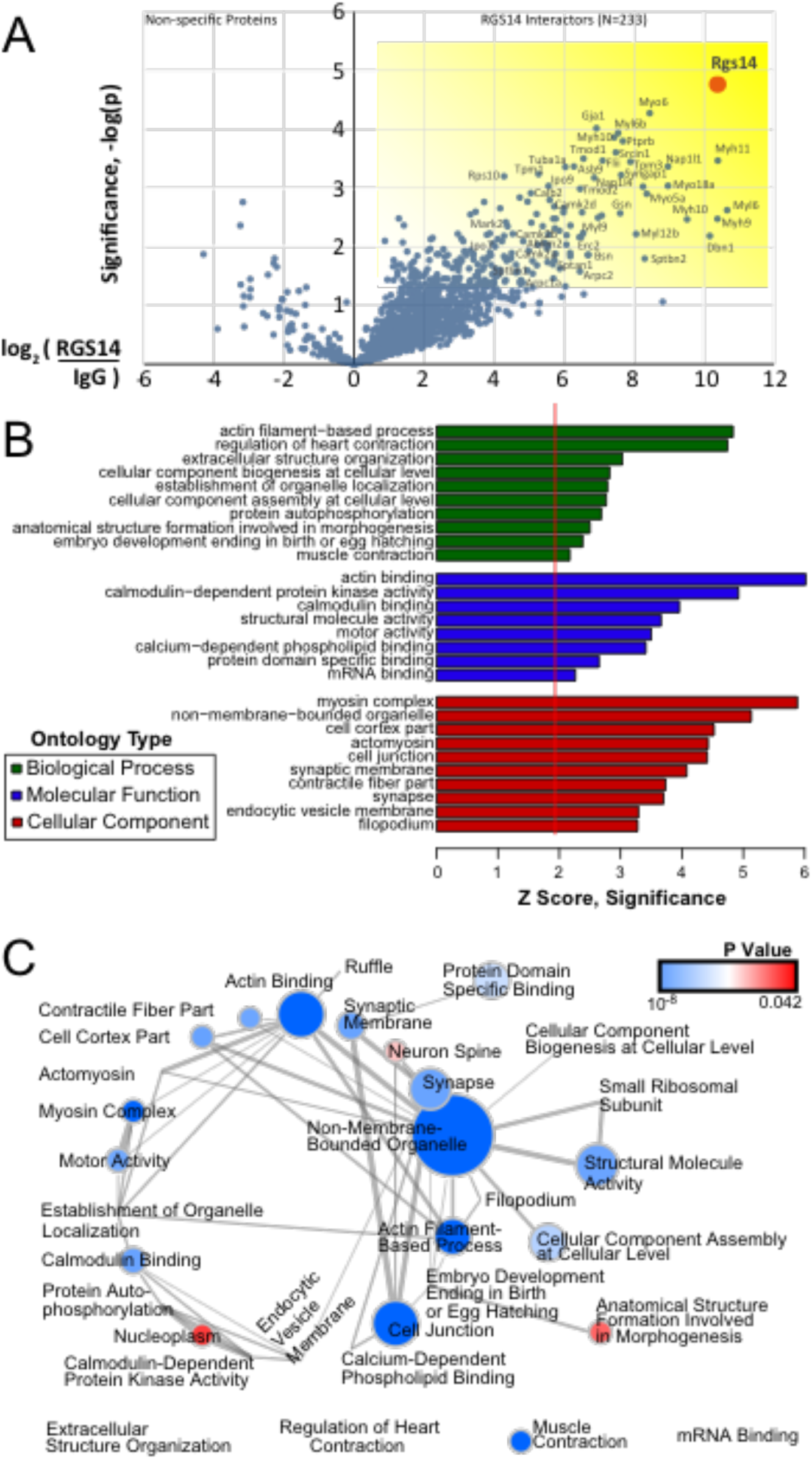
Global analysis of RGS14 interactors from whole mouse brain. (A) Volcano plot of RGS14 interactors (n=3) enriched in pairwise experiments versus nonspecific IgG bead control pulldowns (n=3). A total of 233 specific interactors of RGS14 were determined to be significantly enriched p&0.05 and at least 1.5 fold over the average of the bead controls. (B) Overrepresented gene ontologies among the 233 RGS14 interactors were examined with GO-Elite, and up to the top 10 terms for each of 3 ontology types are shown. Z score of 1.96 represents a significant overrepresentation (p&0.05) relative to the background list of proteins identified in the experiment. (C) Enrichment map of all GO hits is visualized, with nodes representing discrete terms. Node size ranges from invisible (n=5 genes) to large (n=75 genes), and edges correspond to the number of genes overlapping between different terms. Four terms with no overlap to any of the identified terms are shown at the bottom. P value of GO-Elite overrepresentation significance ranged from 6 × 10^−8^ (blue) to 0.042 (red).

**Figure 3.**
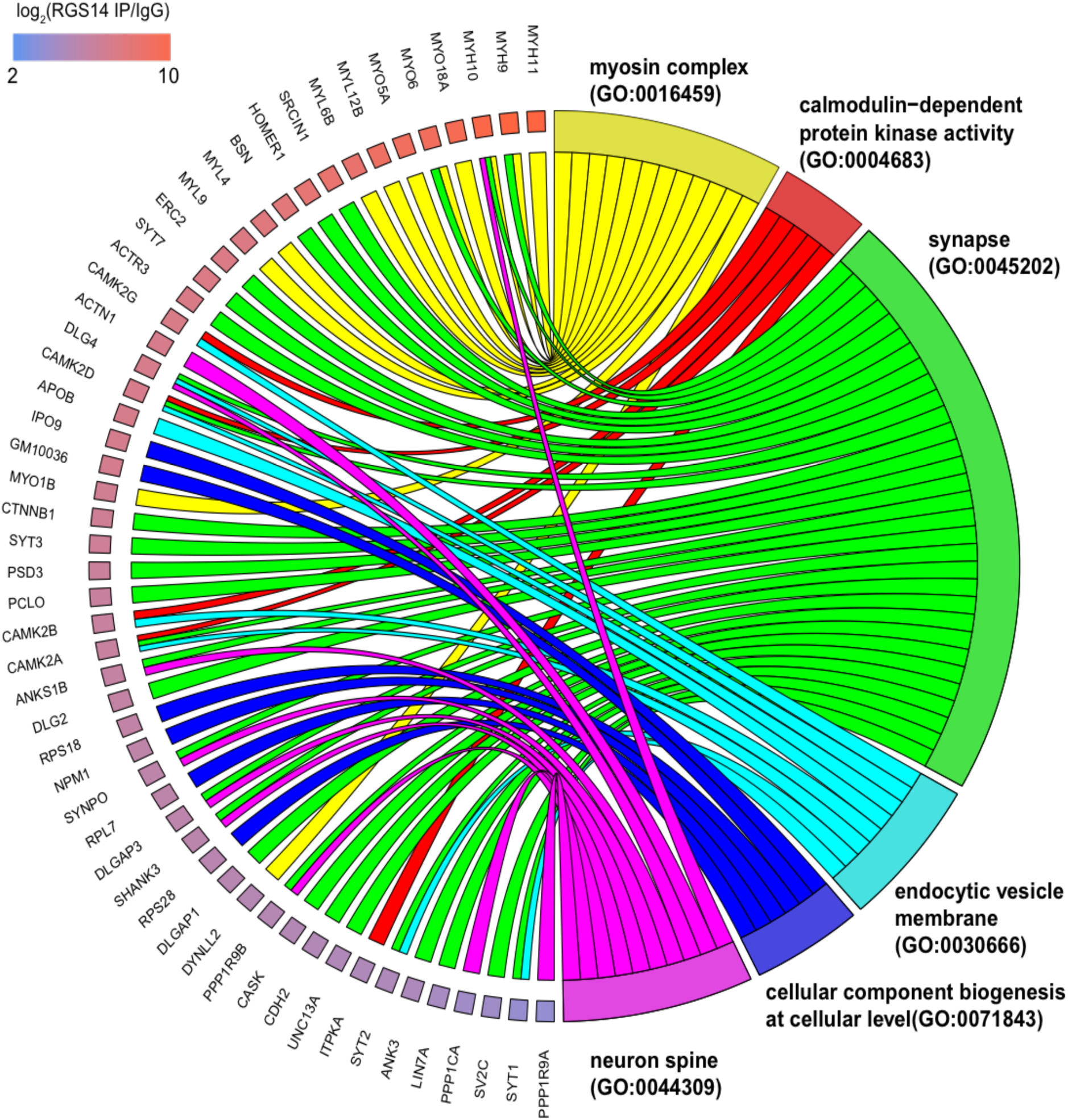
A circular view of RGS14 interactor overlap across selected ontologies. A color scale for individual genes depicts the log_2_(RGS14 IP/IgG) enrichment, which is the sort order for the genes around the semicircle, from highest (top) to lowest (bottom).

Using the CaM target database ^36^, two putative CaM-binding domains were identified in the tandem Ras Binding Domain (RBD) region of RGS14. Performing CaM-Agarose pull-down assays with purified RGS14 protein, we found RGS14 directly interacts with CaM, but only in the presence of Ca^2+^ (Fig 4A). We validated this direct, Ca^2+^ ‐dependent interaction with fluorescence spectroscopy measurements of purified dansyl-CaM and RGS14 observing an increase in dansyl fluorescence (indicative of binding) only in the presence of Ca^2+^ (Fig 4B), revealing a 1:1 binding stoichiometry between Ca^2+^/CaM and RGS14 ^18^. Note that CaM was also identified in RGS14 IP samples by MS analysis, but the enrichment was not significant – likely due to the presence of the Ca^2+^ chelator EDTA in the homogenization buffer (as a necessary protease inhibitor), which disrupts binding of RGS14 to CaM (Fig 4A). To identify the region on RGS14 where CaM binds, we performed CaM-Agarose pull-downs with cell lysates expressing different regions of RGS14 and immunoblotting for FLAG-tags on the truncation mutants (Fig 4C). These experiments confirmed that Ca^2+^/CaM binds RGS14 in the tandem RBD region containing the predicted CaM-binding sites (Fig 4C, asterisks), since only full-length RGS14 and the RBD construct containing the putative CaM binding motifs pulled down with Ca^2+^/CaM. Using differential hydrogen/deuterium exchange (HDX) mass spectrometry we found that Ca^2+^/CaM binding to RGS14 causes increased deuterium incorporation in the tandem RBD region flanking the predicted CaM-binding domains (Figure 4D), and the deuterium incorporation into these peptides becomes significant over time (data not shown). Taken together, these data reveal a previously unknown interaction between RGS14 and Ca^2+^/CaM, and the interaction induces structural changes near predicted CaM-binding domains on RGS14 that are opposite from the decreased deuteration observed with G protein interactions ^10^.

**Figure 4.**
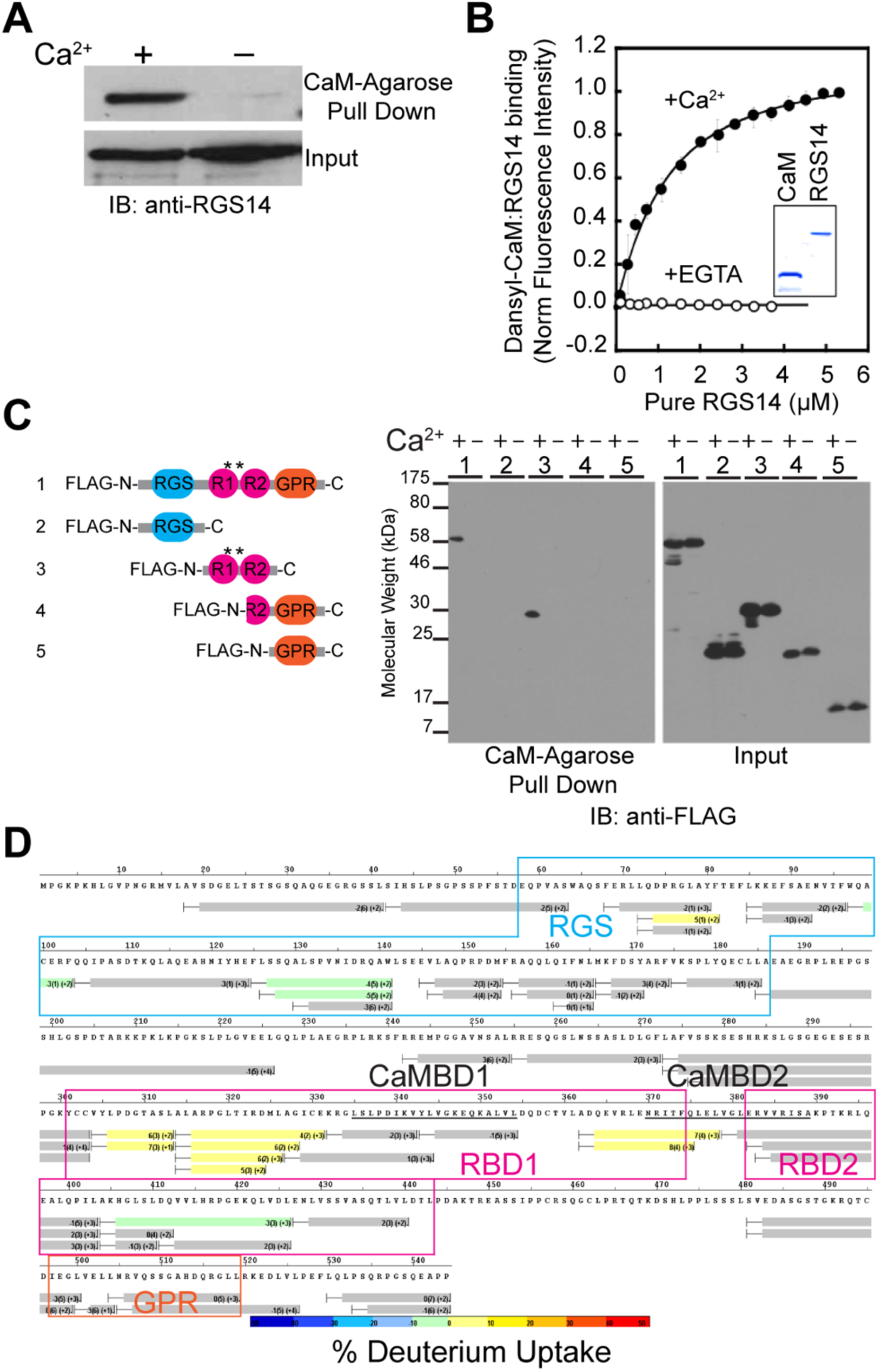
RGS14 directly interacts with Ca^2+^/CaM in the tandem RBD region. A) CaM-Agarose beads pull down purified recombinant RGS14 specifically in the presence of Ca^2+^. No interaction is detected in the presence of a Ca^2+^ chelator (−Ca^2+^). B) Dansyl-CaM fluorescence binding assays confirm that RGS14 directly interacts with Ca^2+^/CaM, but not apo-CaM. Inset: Coomassie stained SDS-PAGE gel of purified recombinant Dansyl-CaM and RGS14 proteins. Data are represented as mean normalized fluorescence intensity ± SEM for three replicates. C) RGS14 interacts with Ca^2+^/CaM through its tandem RBD region. ***Left***: schematic representation of the domain structure of FLAG-tagged RGS14 truncation mutants used to map the interacting region with Ca^2+^/CaM. Asterisks indicate the location of the predicted CaM binding sites. ***Right***: Full-length FLAG-tagged RGS14 or truncation mutant cDNAs expressed in HeLa cells were pulled down by CaM-Agarose beads in the presence or absence of Ca^2+^. Recovered proteins were then subjected to SDS-PAGE and immunoblotting with an anti-FLAG antibody (right). Immunoblotting of input cell lysates with an anti-FLAG antibody verifies expression of all constructs (right). D) A differential HDX heat map for the RGS14•Ca^2+^/CaM complex. Each bar represents an individual peptide with the color corresponding to the average percentage change in deuterium exchange between apo-RGS14 and RGS14•Ca^2+^/CaM over six time points (10, 30, 60, 300, 900, and 3600 s). The numbers in the first parentheses indicate the SD for three replicates. The numbers in the second parentheses indicate the charge of the peptide. Boxed regions indicate residues corresponding to the RGS (blue), RBDs (magenta), and GPR motif (orange); black bars underline the two predicted CaM binding motifs. Peptides are colored according to the color scale bar with percentage of change in deuterium exchange.

We hypothesized the unique conformational changes in RGS14 induced by Ca^2+^/CaM binding may place RGS14 in a favorable conformation to interact with a downstream CaM effector. We focused on CaMKII as a candidate because all four isoforms were were some of the most significantly enriched RGS14 interacting proteins (Fig 2A and 3) and it is required for some forms of CA2 synaptic potentiation ^37^. Co-immunoprecipitation experiments with recombinant RGS14 and CaMKII co-expressed in HeLa cells revealed that RGS14 binds to CaMKII in a Ca^2+^-independent manner (Fig 5A). Further, *in vitro* radiolabeling assays demonstrated that RGS14, but not Gαi1 (negative control), is directly phosphorylated by CaMKII (Fig 5B). These findings indicate that RGS14 interacts with CaMKII in cells independent of Ca2^2+^, and that RGS14 is a CaMKII phosphorylation substrate.

**Figure 5.**
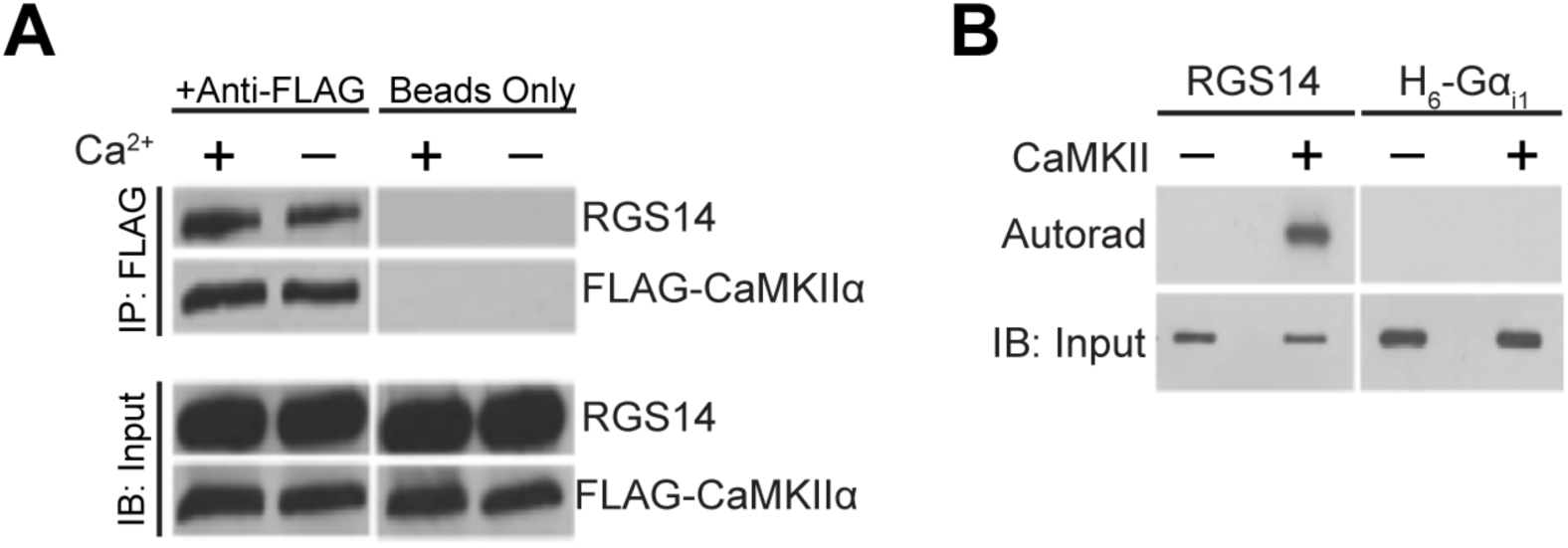
CaMKII directly phosphorylates RGS14 *in vitro*. A) HeLa cells co-transfected with RGS14-Luciferase and FLAG-CaMKIIa cDNAs were lysed, and protein complexes were immunoprecipitated with anti-FLAG agarose beads (“+Anti-FLAG”) or Protein G Sepharose beads (“Beads Only”) as a negative control. Immunoblotting of input cell lysates verifies expression of both cDNAs. B) Purified RGS14 is directly phosphorylated by CaMKIIα *in vitro*. Purified RGS14 or H_6_-Gα_i1_ (negative control) protein were subjected to radiolabeling assays with ^32^P-ATP in the presence/absence of purified CaMKIIα, and phosphorylation was detected by SDS-PAGE and subsequent autoradiography. Immunoblotting was performed on input proteins to verify protein levels.

To determine if endogenous RGS14 interacts with CaM and CaMKII in hippocampal CA2 neurons, we performed proximity ligation assays (PLA) in mouse brain sections. This technique allows visualization of protein interactions in fixed tissue by producing red fluorescence only when two immunolabeled proteins are within close proximity (approximately 40 nm) ^33^. Interactions between RGS14 and CaM were weakly visible in area CA2 of mouse hippocampus (Fig 6A,C), albeit at higher intensity levels compared with fluorescence in area CA1 (negative control) in the same sections (Fig 6 B,D). By contrast, RGS14 interactions with CaMKII were clearly visible in CA2, but also in CA1 at lower intensity levels. The latter data can be explained by the fact that CA2 pyramidal neurons with RGS14-containing axons project to area CA1 (Fig 1A), or alternatively, by RGS14 protein expression we spuriously observe in CA1 pyramidal cells ^2^. These results indicate that RGS14 interacts with its newly validated binding partners in hippocampal area CA2, and further suggest a previously unrecognized role for RGS14 in regulating key Ca^2+^ signaling pathways required for plasticity in CA2 neurons.

**Figure 6.**
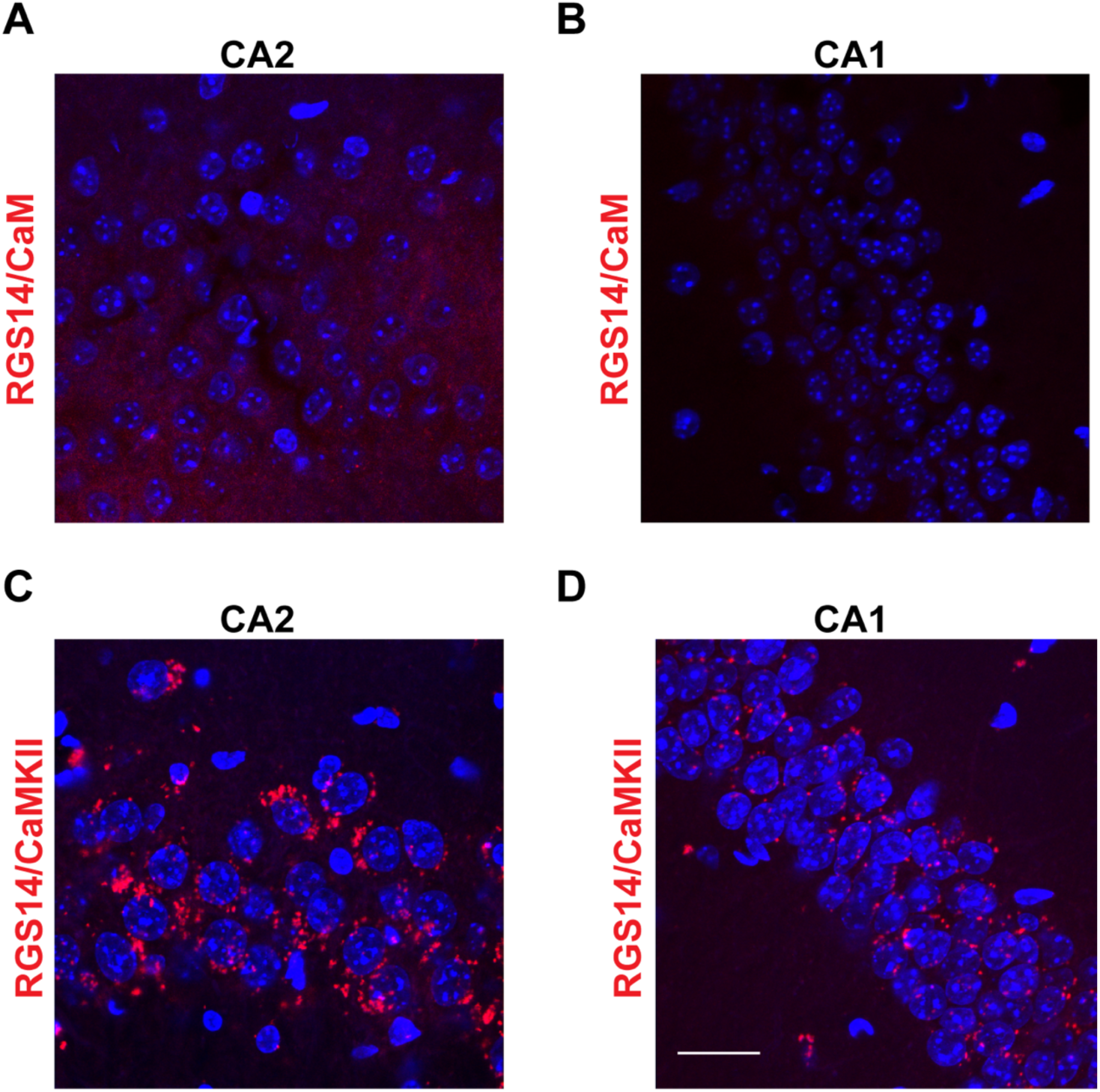
RGS14 interacts with endogenous CaM and CaMKII in mouse hippocampal CA2 neurons. Confocal images of hippocampus sections following proximity ligation assays to produce protein interaction-dependent fluorescence (red) between RGS14 and CaM (A,B) or RGS14 and CaMKII (C,D). Both interactions are markedly higher in hippocampal area CA2 (A,C) compared with area CA1 (B,D; negative control). Images are representative results of three independent replicates. Scale bar = 25 μm.

## Discussion

In this study, we find that RGS14 exists in one or more high molecular weight protein complexes in mouse brain tissue and identify the constellation of proteins (n = 233) that interact with endogenous RGS14 in brain. Further, we validate and characterize direct interactions between RGS14 and two novel binding partners revealed by this proteomic study. We find that RGS14 directly binds to Ca^2+^/CaM within the tandem RBD region, which results in destabilization (i.e. increased deuterium incorporation) in the same region. In contrast, RGS14 binds to CaMKII in a Ca^2+^-independent manner in cells and is directly phosphorylated by CaMKII *in vitro*. Lastly, we show that RGS14 interacts with both CaM and CaMKII in its host CA2 neurons in hippocampus providing support for the physiological relevance of these findings.

These studies are the first report to investigate the network of signaling proteins that RGS14 natively engages in mouse brain, either directly or indirectly, to suppress neuronal plasticity and learning and memory. Obviously, RGS14 does not directly interact with all 233 of these proteins, but likely interacts with key node proteins that form a complex with other functionally related proteins. However, our results do identify two new RGS14 direct binding partners – Ca^2+^/CaM and CaMKII – that have been previously implicated in the suppression ^35^ and expression ^37^ of plasticity in CA2 neurons, respectively. Our findings also suggest a novel role for RGS14 in regulating key postsynaptic Ca^2+^-driven pathways required for plasticity induction, providing a new potential mechanism upstream of Ras/ERK by which RGS14 could gate synaptic plasticity. This is the first study to link RGS14 to Ca^2+^ regulation in neurons and is consistent with a previous report showing robust Ca^2+^ handling in CA2 neurons contributes to limited plasticity therein ^35^. Future studies will investigate whether RGS14 regulates spine Ca^2+^ transients and downstream signaling in CA2 neurons.

The list of candidate RGS14 interacting proteins identified here suggests exciting new possible mechanism(s) by which RGS14 could restrict synaptic plasticity. The most enriched proteins (Fig 3) have a range of functions in postsynaptic mechanisms of LTP including: 1) Ca^2+^-activated signaling molecules, 2) cytoskeletal molecules (actin/myosin binding) important for AMPA-type glutamate receptor trafficking and structural rearrangements, 3) protein kinases/phosphatases that regulate phosphorylation state of synaptic targets, 4) scaffolding proteins that assemble and anchor signaling complexes within spines. Future studies will explore functional interactions between RGS14 and these proteins. The wide array of proteins that interact with RGS14 in brain may suggest that multiple signaling pathways converge on RGS14 allowing it to serve as a signaling platform that suppresses plasticity through multiple mechanisms. Alternatively, a few key proteins, such as Ca^2+^/CaM or other scaffolding proteins, could serve as signal integration platforms to recruit and allow RGS14 to modulate the activity of the proteins they assemble. While most of the protein partners for RGS14 we identified play key roles in postsynaptic plasticity, consistent with RGS14 roles in suppressing LTP, we note that RGS14 is also present in CA2 axons that project to CA1 ^1,3^. Consistent with this finding, we have recently observed that endogenous RGS14 is present at CA2 terminals that synapse onto neurons of CA1 in monkey ^3^. Here we observe notable interactions between RGS14 and CaMKII in CA1, either due to postsynaptic expression of RGS14 in CA1 somata and dendrites or presynaptically within CA2 nerve terminals, though presynaptic functions of RGS14 in CA2 axons remain unexplored.

Noticeably absent from the list of prominent RGS14 binding partners were the previously demonstrated signaling partners, H-Ras and Rap2 (note that some minor amount of Gαi/o binding was observed). A likely explanation for this observation is that the binding of each of these proteins to RGS14 requires an activation step which would be largely absent in unstimulated brain homogenates. By contrast, the newly identified RGS14 binding partners, in particular the actomyosin cytoskeletal binding proteins, suggests that RGS14 may play previously unappreciated roles in regulating spine plasticity, and that perhaps H-Ras/Rap2 and Gαi/o-GTP binding only occurs transiently in response to a stimulus and activation step, as we have shown previously in cultured cells ^10^. In this model, GTPase binding could modulate RGS14 regulation of spine morphology following postsynaptic excitatory input. Alternatively, RGS14 may predominantly interact with G protein and Ras/Rap binding partners in non-neuronal cells, i.e. in heart or immune cells ^9^. With our new findings here, we now can add Ca^2+^/CaM binding and CaMKII phosphorylation, either alone or in some combination with a G protein and H-Ras/Rap2 binding, as possible regulatory components of RGS14 actions in spines. Future studies will explore these possibilities.

The question still remains as to why RGS14 would exist to naturally suppress plasticity in CA2 neurons and hippocampus-dependent learning. Strong evidence supports a role for CA2 neurons in encoding spatial, temporal, and social aspects of memory ^1,38–41^. A previous study of the spatiotemporal expression pattern of RGS14 in the mouse brain found that RGS14 mRNA/protein levels are absent at birth, whereas both are gradually upregulated throughout postnatal development until reaching highest levels in adult brain ^2^. We hypothesize this developmental regulation of RGS14 expression could allow for a period of enhanced plasticity and learning in early adolescence when spatial navigation and social recognition memory are critical for survival, and later RGS14 expression would allow for regulated plasticity as an information filter into hippocampal pathways important for learning and memory. Future studies are necessary to determine if RGS14 regulates social behavior and temporal order of events.

## Supporting Information

Table of complete spectra of MaxQuant protein groups output (Supp. Table 1); table of enrichment statistics for proteins identified in IP samples (Supp. Table 2).

## Acknowledgments

P.R.E. was supported by a predoctoral fellowship from the National Institutes of Health/National Institute of Neurological Disorders and Stroke (1F31NS086174). J.R.H. was supported by grants from the National Institutes of Health/National Institute of Neurological Disorders and Stroke (5R01NS37112 and 1R21NS074975). S.M.D. was supported by the Intramural Research Program of the National Institute of Environmental Health Sciences, National Institutes of Health (Z01ES100221). Research reported in this publication was supported in part by the Emory Neuroscience NINDS Core Facilities and NIH/NINDS under award number P30NS055077.

